# TEFAR: a lightweight, configurable framework for semi-automatic artefact-component classification in EEG and TMS–EEG

**DOI:** 10.64898/2026.09.22.751113

**Authors:** Christina Dimitriadou, Nicholas Furl

**Affiliations:** Department of Psychology, Royal Holloway, University of London, Egham TW20 0EX, United Kingdom

**Keywords:** electroencephalography, transcranial magnetic stimulation, independent component analysis, artefact rejection, FieldTrip, preprocessing, open-source software

## Abstract

Independent component analysis (ICA) is widely used to remove artefacts from EEG and TMS–EEG recordings, but the components that correspond to artefacts are usually identified by manual inspection, which is subjective and difficult to reproduce. Existing automated classifiers are either designed for ordinary EEG and do not target the artefacts specific to concurrent transcranial magnetic stimulation, or are built on EEGLAB, so none is available for FieldTrip-based pipelines. We present TEFAR, a lightweight, configurable framework for semi-automatic artefact-component classification that applies the same detectors to ordinary EEG and TMS–EEG. TEFAR is built on FieldTrip and requires no additional MATLAB toolboxes. Each artefact class (line noise, blinks, lateral eye movements, cranial muscle, cardiac activity, and the TMS decay and recharge artefacts) is identified from the spectral, spatial, or temporal signature that its components are known to express. Detection thresholds use robust statistics (the median and the median absolute deviation rather than the mean and standard deviation), so that a single dominant artefact component cannot raise the threshold that is meant to detect it. In 30 ground-truth simulations, TEFAR reached a specificity of 1.000 and a sensitivity of 0.96–0.97 for both profiles. An application of TEFAR to real EEG and TMS–EEG recordings is the next step and will be reported separately.

## 1 Introduction / Statement of need

Independent component analysis (ICA) is the standard method for removing stereotyped artefacts (ocular, muscular, cardiac, and line noise) from electroencephalographic (EEG) recordings (Delorme & Makeig, 2004; Hyvärinen & Oja, 2000). After the decomposition, the components that correspond to artefacts must still be identified and removed, and in most laboratories this is done by manual inspection. Manual selection is time-consuming, requires expertise, and is difficult to reproduce across raters and studies. For this reason, several automated component classifiers have been developed (Chaumon et al., 2015).

These include ADJUST (Mognon et al., 2011), FASTER (Nolan et al., 2010), MARA (Winkler et al., 2011), SASICA (Chaumon et al., 2015), and the machine-learning classifier ICLabel (Pion-Tonachini et al., 2019). These classifiers have substantially improved the reproducibility of artefact removal, but they have two limitations for the present use case. First, most of them are implemented within EEGLAB (Delorme & Makeig, 2004) and depend on additional resources (extra MATLAB toolboxes or, in the case of ICLabel, a large training corpus). As a result, no lightweight classifier is available for FieldTrip-based pipelines. Second, they were developed for and validated on ordinary EEG, and they do not target the artefacts that are specific to concurrent transcranial magnetic stimulation and EEG (TMS–EEG).

TMS–EEG recordings contain artefacts that are absent from ordinary EEG (Rogasch et al., 2014, 2017). These include the large decay artefact (a step or exponential deflection) that follows the pulse, the later recharge artefact of the stimulator, and focal cranial-muscle components evoked near the coil. Removing these artefacts while preserving the neural response is a central methodological problem in TMS–EEG, and the general-purpose classifiers listed above do not address it. The TESA toolbox (Rogasch et al., 2017) provides semi-automatic component classification for TMS–EEG, but it operates within EEGLAB. Other TMS–EEG tools, such as ARTIST (Wu et al., 2018) and TMSEEG (Atluri et al., 2016), also operate within EEGLAB. No comparable classifier exists for FieldTrip-based pipelines, and none applies the same detectors to ordinary EEG and TMS–EEG.

Here we introduce TEFAR (TMS-EEG / EEG FieldTrip Artefact Removal), a lightweight tool for semi-automatic artefact-component classification in ordinary EEG and TMS–EEG. TEFAR is built on FieldTrip (Oostenveld et al., 2011) and requires no additional MATLAB toolboxes; the statistical, spectral, and temporal features that it uses are implemented internally. TEFAR consists of three MATLAB functions: a core function, which runs ICA and classifies the components, and two profile functions, one for ordinary EEG and one for TMS– EEG, which set the defaults of each profile and call the core function. All settings are fields of a MATLAB structure array, `cfg`, which the user can pass as an optional input, following the FieldTrip convention. Each artefact class is detected from an established independent-component signature. Each detector computes one feature value per independent component and flags the components whose value is extreme relative to the other components. The detection thresholds are expressed as robust scores, based on the median and the median absolute deviation (MAD) rather than on the mean and standard deviation. The reason is that a successful decomposition typically concentrates an artefact in a single component, so the feature value of this component is an outlier among the values of all components. This outlier increases the mean and the standard deviation on which a conventional threshold is based, and the artefact component may then not exceed the threshold. The median and the MAD are almost unaffected by a single outlier (see Design, Robust thresholds).

TEFAR is semi-automatic by design. It suggests components for removal, but the final decision rests with the researcher. Visual inspection of component topographies, time courses, and spectra remains a required step of the workflow. This is current best practice in TMS– EEG (Rogasch et al., 2017). It is also necessary because the quality of the ICA decomposition varies across datasets (Ryali et al., 2009; Tamburro et al., 2021), and because artefactual and neural activity can overlap closely in time after the pulse (Atti et al., 2024). TEFAR is therefore intended to accelerate and standardise component selection, not to replace expert judgement.

## 2 Design

### Architecture

TEFAR consists of a core function, `tefar_core`, and two profile functions, `TEFAR_eeg` and `TEFAR_tms`. The user calls the profile function for the data type, with the FieldTrip data structure as input and, optionally, a `cfg` structure. The profile function sets the defaults of its profile for every field that the user did not specify, and passes `cfg` and the data to `tefar_core`. The core function performs ICA with the FastICA algorithm (Hyvärinen & Oja, 2000) through FieldTrip’s `ft_componentanalysis` (or accepts a precomputed component structure). For every independent component, it computes a set of features and applies the detectors. It returns the FieldTrip component structure and a results structure with the indices of the components flagged by each detector. The two profile functions differ only in their defaults: the EEG profile enables the line, blink, lateral eye-movement, muscle, and cardiac detectors; the TMS profile enables the line, blink, muscle, and cardiac detectors, and adds the decay and recharge detectors, which are specific to TMS–EEG.

Every frequency band, threshold, temporal window, and channel set is a field of `cfg` with a documented default that can be overridden. Adapting TEFAR to a new montage, sampling rate, or mains frequency (for example, 60 Hz) therefore requires only a field assignment, not a change to the code.

### Detectors

Each artefact class is identified from an established independent-component signature (Table 1). These signatures are well documented in the artefact literature, and several of them are already used by existing classifiers: the spatial blink and lateral eye-movement features of ADJUST (Mognon et al., 2011), the focal-topography measure of Chaumon et al. (2015), and the post-pulse amplitude criteria of TESA (Rogasch et al., 2017). TEFAR combines these rules in a single configurable core function and expresses their thresholds in robust (median/MAD) form.

**Table 1.**
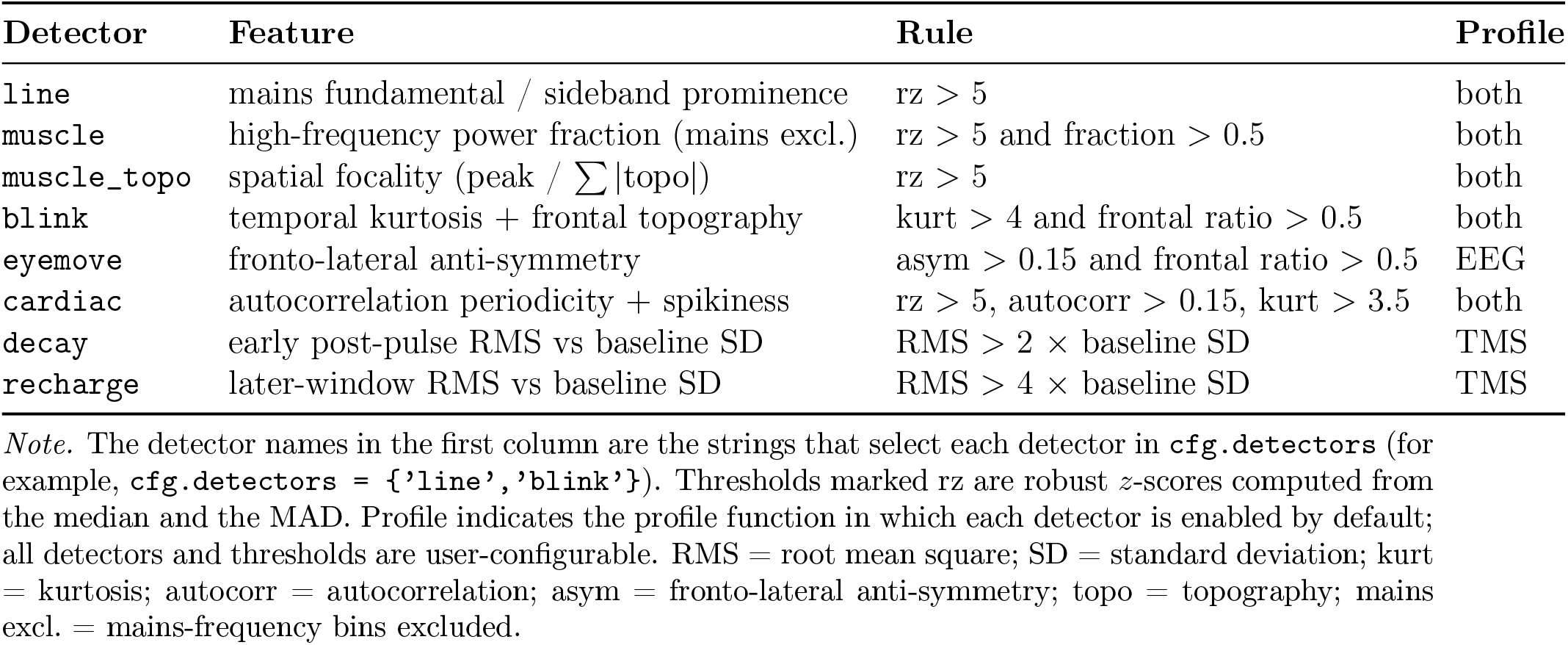
Artefact detectors and their discriminating features.

Spectral features are derived from a multitaper power spectrum. Line noise is detected as the prominence of the mains fundamental relative to its spectral sidebands; cranial muscle activity (EMG) as the fraction of power in a high-frequency band, with the mains bins excluded (Goncharova et al., 2003). Spatial features are derived from the component topography. Focal components (muscle and the TMS decay and recharge artefacts) are flagged by high spatial focality, defined as the ratio of the peak absolute weight to the summed absolute weight (Chaumon et al., 2015). Ocular components are identified from their frontal concentration; lateral eye movements are further distinguished from blinks by an anti-symmetric left–right frontal weighting (Mognon et al., 2011). The remaining classes are identified from temporal features. Blinks are flagged by the high kurtosis of the component time course (Delorme et al., 2007); cardiac components by a periodic autocorrelation peak at a physiological lag, combined with spikiness. The TMS decay and recharge artefacts (Veniero et al., 2009) are detected from their root-mean-square (RMS) amplitude in defined post-pulse windows, relative to the pre-stimulus baseline. The temporal windows are defined in seconds relative to the TMS pulse (time zero). In each trial, the samples inside a window are selected from the time axis of that trial, not from fixed sample indices. The detectors therefore make no assumption about the number or length of the trials.

### Robust thresholds

A design choice with a large effect on performance is the use of robust (median/MAD) *z*-scores instead of the conventional mean + *k* SD rule (Leys et al., 2013). Under the conventional rule, an independent component from the ICA is flagged when its feature value (for example, its kurtosis or its high-frequency power fraction) exceeds the mean of that feature across all components by *k* standard deviations. Each feature in Table 1 is computed for every independent component, so a detector operates on a distribution of *N* values, one per component. When a single component carries almost all of an artefact (as is typical for line noise or a large muscle component), its extreme value increases both the mean and the standard deviation of this distribution. The conventional criterion can then no longer be met: for one outlier among *N* components, the maximum attainable *z*-score is 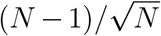, approximately 4.0 for *N* = 18. A threshold of four standard deviations is therefore equal to the maximum that even an arbitrarily large artefact can produce. Replacing the mean and standard deviation with the median and the MAD removes this dependence on the outlier itself, and the resulting thresholds remain stable across widely varying numbers of components. Two thresholds are absolute rather than robust. Kurtosis is scale-free and equals 3 for a Gaussian process, so the blink criterion of 4 identifies clearly supra-Gaussian activity. The cardiac spikiness criterion of 3.5 excludes sinusoidal components, for which kurtosis is exactly 1.5.

### Semi-automatic operation

TEFAR reports the components flagged by each detector, together with a per-component score (the number of detectors that flagged that component as an artefact). Components are suggested for removal either as the union of all detectors (the default) or when at least a configurable number of detectors agree. In keeping with best practice for TMS–EEG (Rogasch et al., 2017), TEFAR is intended to accelerate and standardise a decision that the researcher confirms by visual inspection; it does not remove components automatically.

## 3 Validation

### Ground-truth simulation

Because the artefact identity of a real independent component is never known with certainty, we validated TEFAR on simulated data generated under the ICA model, **x** = **As**. Each latent source is assigned a known label and a known scalp projection (a column of the mixing matrix **A**); the projections are specified directly at the electrode level and are not derived from a dipole forward model. Every component that TEFAR classifies can therefore be scored against the known label. Brain sources are modelled as 1*/f* background activity with an oscillatory peak in the theta–alpha range and random topographies. The injected artefacts reproduce the canonical signatures: a mains sinusoid with a broad topography; sparse frontal blink deflections; a broadband high-frequency muscle source with a focal lateral topography; quasi-periodic cardiac spikes; and, for the TMS profile, an early exponential decay artefact and a later recharge artefact, both with focal topographies. All artefacts of a profile are present in every simulated dataset, each as a separate source with a fixed amplitude, so each component carries exactly one artefact class or brain activity. Unless otherwise noted, each simulation comprised twelve brain sources and the artefact set of the profile. These sources were projected onto a 30-channel 10–20 montage, with 40 epochs sampled at 1 kHz. The simulation depends only on base MATLAB and regenerates every source, topography, and mixing matrix from a fixed seed. It therefore runs without FieldTrip and is a self-contained, reproducible benchmark.

### Evaluation

We evaluated detection at the component level. In this mode, the simulated sources (their time courses and topographies) are passed to TEFAR in place of the output of ICA, and TEFAR classifies them without access to their labels. The labels are used only afterwards, to score each decision as correct or incorrect. Because ICA is not run, any error is due to the detectors and not to an imperfect separation of the sources by ICA. An end-to-end mode is also provided: it mixes the sources to channels, runs FastICA, matches each recovered component to the true source with the most similar topography, and then scores the decisions in the same way. Performance was quantified as the sensitivity, specificity, and accuracy of the artefact-versus-brain decision. Figure 1 shows that each simulated artefact reproduces its expected signature in topography, spectrum, and time course. Figure 2 shows the corresponding feature values across components: for every detector, the target artefact exceeds the decision threshold and the brain components remain well below it. Two detectors (blink and cardiac) combine a primary feature with a second condition: frontal topography and spikiness, respectively. A component such as the decay artefact, despite its high kurtosis, is therefore correctly not classified as a blink (its topography is not frontal) or as cardiac (it is not periodic).

**Figure 1.**
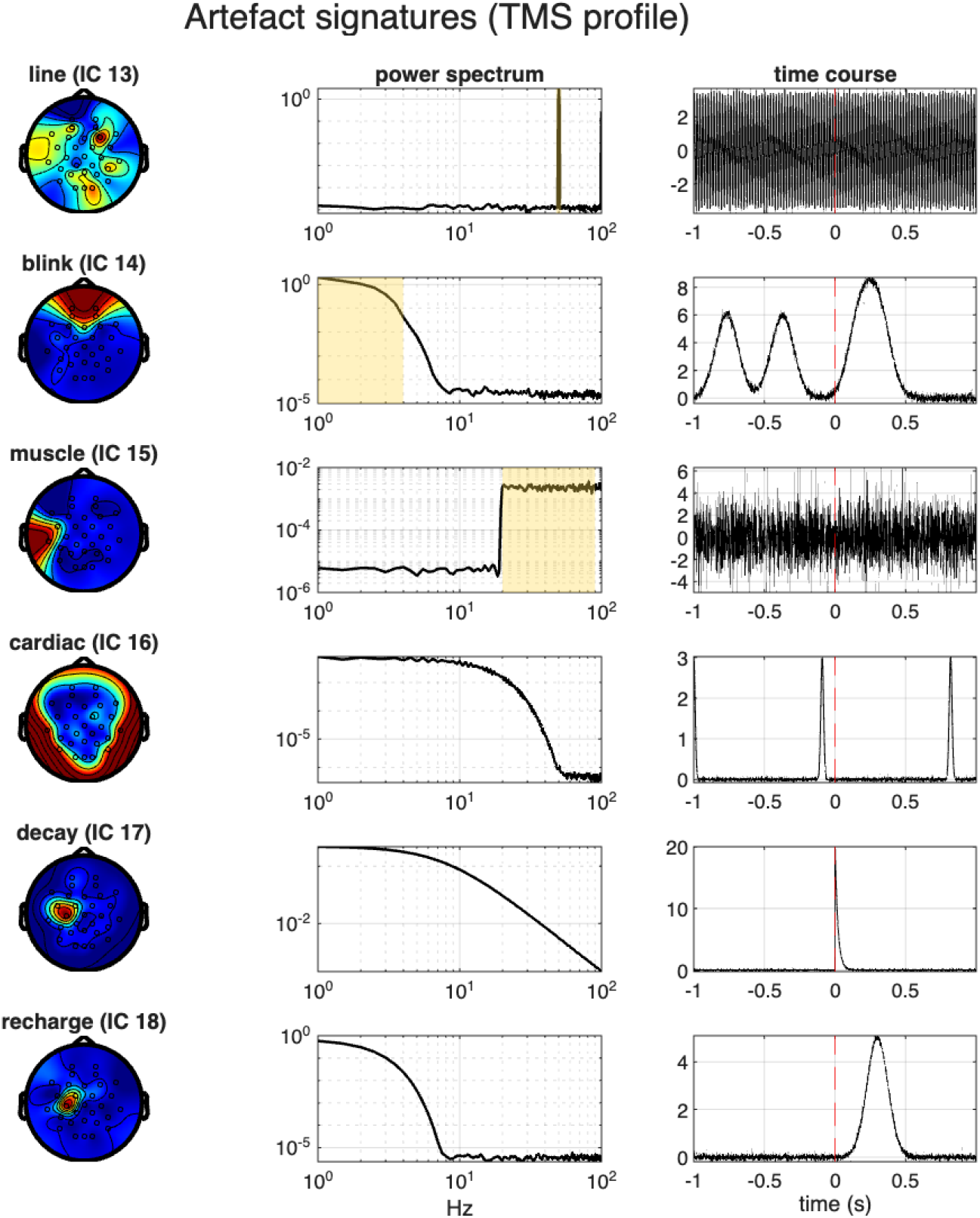
Artefact signatures (TMS profile). Each row shows the topography, the power spectrum (diagnostic band shaded), and an example time course of one injected artefact.

**Figure 2.**
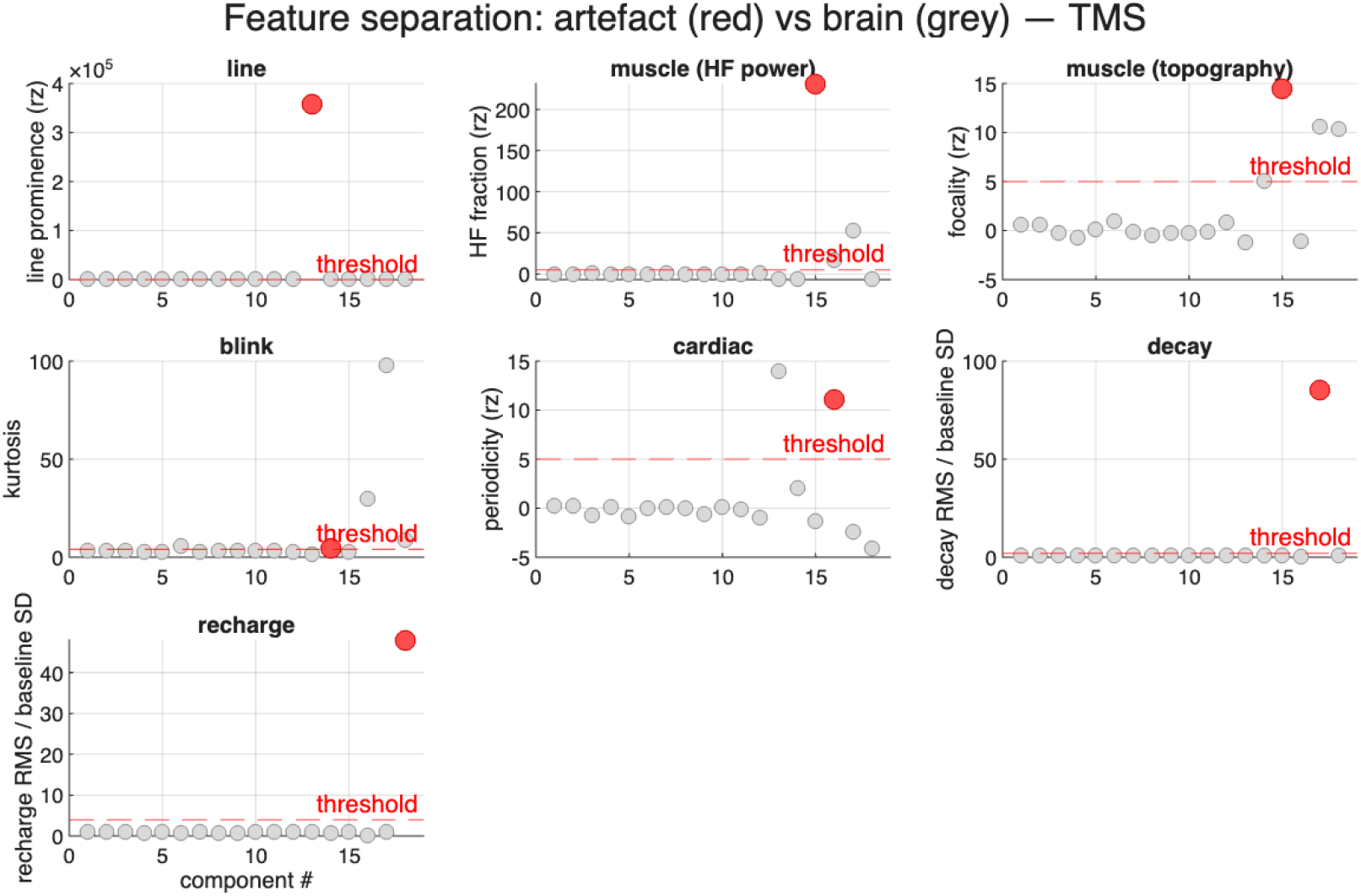
Feature separation (TMS profile). Data are from one simulated dataset with a fixed seed: twelve brain sources and six artefact sources, 40 epochs. Each panel shows one detector. Each point is one component; the metric is computed across all 40 epochs. Red: the component carrying the target artefact. Grey: all remaining components, brain and other artefact classes. The dashed line is the decision threshold. Blink and cardiac also require a second condition (frontal topography and spikiness, respectively), so a grey point above a single threshold is not a false positive.

Across 30 independent simulations, the TMS profile achieved a mean sensitivity of 0.956, a specificity of 1.000, and an accuracy of 0.985 (Figure 3), and the EEG profile 0.973, 1.000, and 0.992, respectively (Figure 6). Specificity is the property that matters most in practice, because a false positive removes brain activity. In these 30 simulations, no brain component was flagged for removal in either profile. The missed artefacts (which account for the lower tail of the sensitivity distribution) were most often blinks whose kurtosis fell just below the threshold. Such borderline cases are the reason for the visual inspection step. To check that 30 seeds were sufficient, we repeated the evaluation with 40 seeds; each seed regenerates every source, topography, mixing matrix, and event timing. Each mean changed by less than 0.01 (TMS: sensitivity 0.962, specificity 1.000, accuracy 0.987; EEG: 0.965, 0.998, 0.988), so the estimates are stable. The EEG specificity of 0.998 corresponds to one false positive among 480 brain components (twelve brain sources in each of 40 seeds). False positives are therefore rare but not impossible, which is a further reason for the visual inspection step.

**Figure 3.**
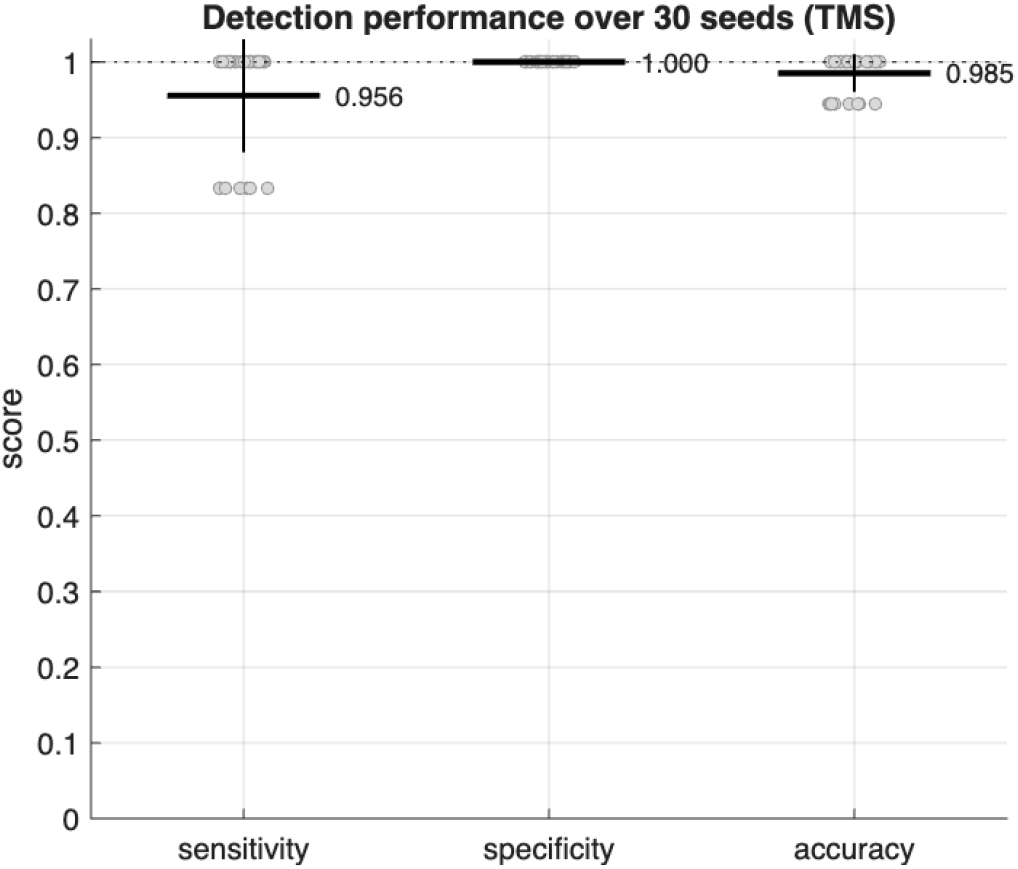
Detection performance over 30 simulated datasets (TMS profile). Points are per-seed scores; bars show mean *±* SD. Specificity was 1.000 in every seed.

The artefact signatures, feature separation, and detection performance of the EEG profile are shown in Figures 4–6. The lateral eye-movement (`eyemove`) detector distinguishes saccadic components from blinks by their fronto-lateral anti-symmetry; the blink detector alone does not separate these two classes.

**Figure 4.**
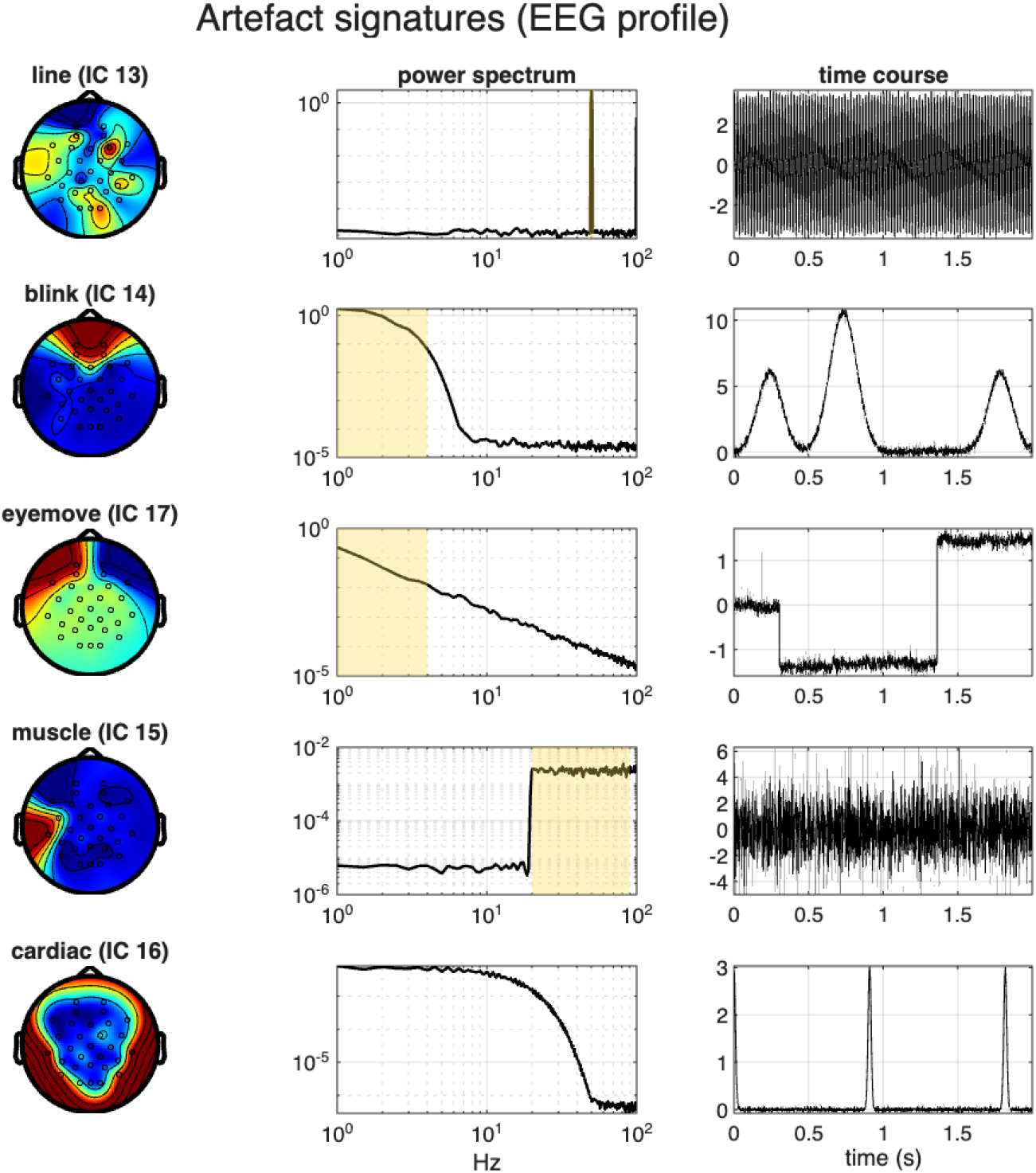
Artefact signatures (EEG profile): topography, power spectrum, and time course of each injected artefact (line, blink, eye movement, muscle, cardiac).

**Figure 5.**
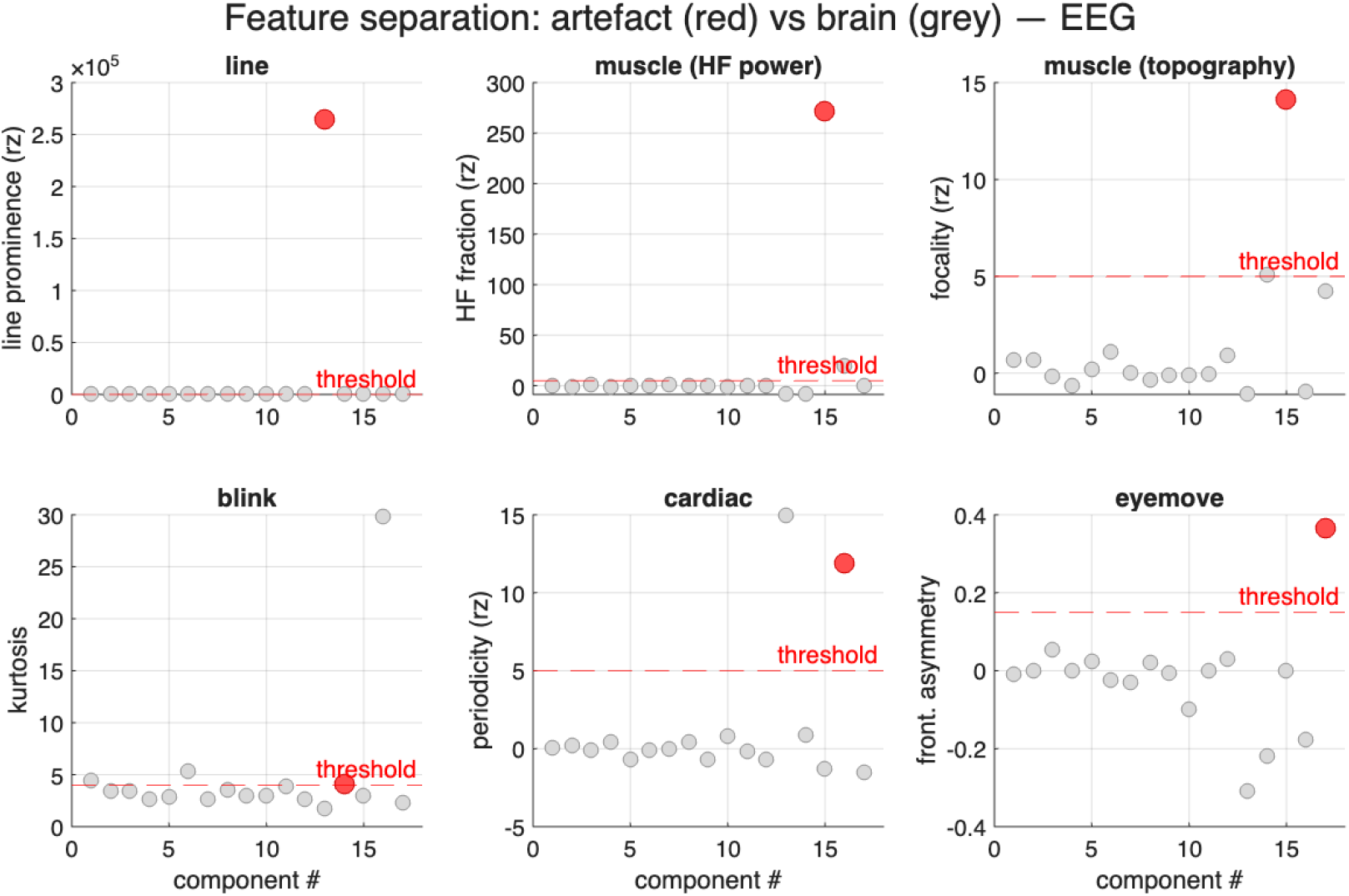
Feature separation (EEG profile). Data are from one simulated dataset with a fixed seed. Each panel shows one detector; each point is one component. Red: the component carrying the target artefact. Grey: all remaining components, brain and other artefact classes. The dashed line is the decision threshold.

**Figure 6.**
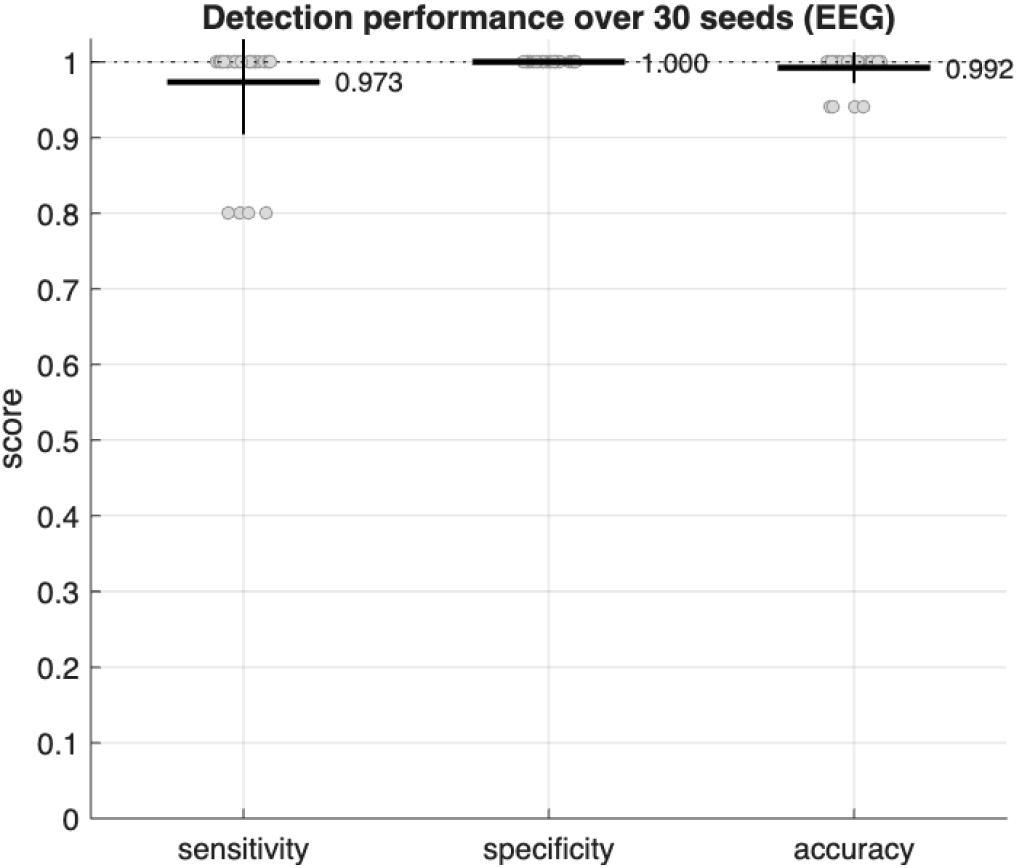
Detection performance over 30 simulated datasets (EEG profile). Points are per-seed scores; bars show mean *±* SD.

## 4 Availability and usage

TEFAR is released as open-source MATLAB code at https://github.com/christinadelta/tefar, under the MIT licence. A tagged release (v1.0.0) is archived on Zenodo (DOI: `10.5281/zenodo.22706419`). The only requirement is a working MATLAB installation with FieldTrip (Oostenveld et al., 2011) on the path; no additional MATLAB toolboxes are needed.

The two profile functions are called as follows:

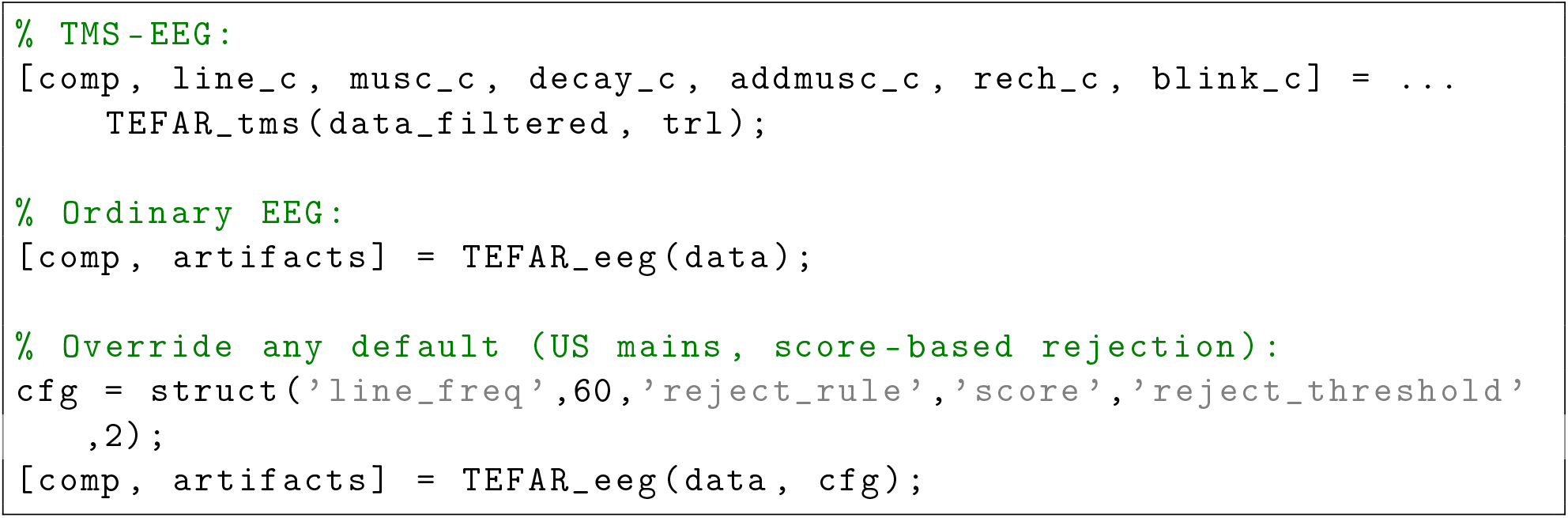

The third example overrides the mains frequency and the rejection rule through `cfg`. The complete validation reported here can be reproduced with the accompanying scripts: `verify_tefar_logic` reproduces the detector results without FieldTrip; `validate_tefar` runs the full pipeline on simulated data; and `make_tefar_figures` regenerates the figures in this paper.

## 5 Limitations

Several limitations should be noted. First, the validation rests on simulated data. Source parameters, topographies, and event timings are randomised across seeds, and measurement noise is added, but the simulated sources express their canonical signatures by construction. The simulation therefore tests whether the detectors and their thresholds behave as intended; it does not test how difficult real data are, and real artefacts need not match these signatures. The simulated data also satisfy the assumptions of ICA exactly (a linear, instantaneous mixture of independent sources), so FastICA is expected to separate them more cleanly than real recordings, which do not satisfy these assumptions. Artefact amplitudes are fixed, so the simulation does not show at which amplitude, or at which noise level, each detector fails; a parameter sweep would establish these limits. Each simulated component carries a single artefact class, whereas in real recordings ICA can leave two classes in one component (for example, the TMS decay artefact together with cranial muscle activity near the coil); the detectors have not been tested on such mixed components. In addition, the simulation is not a physiological forward model; no head geometry or volume conduction is represented. Artefact topographies are constructed by assigning weights to the electrode sites at which each artefact class typically appears (for example, frontal sites for blinks and sites near the coil for the TMS artefacts), not by projecting dipolar sources through a leadfield. For these reasons, the simulation is not a substitute for evaluation on real EEG and TMS–EEG recordings, which we regard as the essential next step before routine use. Second, TEFAR is semi-automatic by design; it suggests components for removal but does not replace expert visual inspection, and it should not be used to remove components unattended. Third, among the detectors, the blink detector is the most marginal, with target kurtosis values close to the threshold; it accounts for most of the missed artefacts in our simulations and should be refined first.

## Data and code availability

The data and code used in this study are available in the GitHub repository tefar.

## Declaration of competing interest

The authors have declared no competing interest.

## Notes

https://doi.org/10.5281/zenodo.21844563

https://github.com/christinadelta/tefar

## References

Atluri, S., Frehlich, M., Mei, Y., Garcia Dominguez, L., Rogasch, N. C., Wong, W., Daskalakis, Z. J., & Farzan, F. (2016). TMSEEG: A MATLAB-based graphical user interface for processing electrophysiological signals during transcranial magnetic stimulation. Frontiers in Neural Circuits, 10, 78. 10.3389/fncir.2016.00078

Atti, I., Belardinelli, P., Ilmoniemi, R. J., & Metsomaa, J. (2024). Measuring the accuracy of ICA-based artifact removal from TMS-evoked potentials. Brain Stimulation, 17 (1), 10–18. 10.1016/j.brs.2023.12.001

Chaumon, M., Bishop, D. V. M., & Busch, N. A. (2015). A practical guide to the selection of independent components of the electroencephalogram for artifact rejection. Journal of Neuroscience Methods, 250, 47–63. 10.1016/j.jneumeth.2015.02.025

Delorme, A., & Makeig, S. (2004). EEGLAB: An open source toolbox for analysis of single-trial EEG dynamics including independent component analysis. Journal of Neuroscience Methods, 134 (1), 9–21. 10.1016/j.jneumeth.2003.10.009

Delorme, A., Sejnowski, T., & Makeig, S. (2007). Enhanced detection of artifacts in EEG data using higher-order statistics and independent component analysis. NeuroImage, 34 (4), 1443–1449. 10.1016/j.neuroimage.2006.11.004

Goncharova, I. I., McFarland, D. J., Vaughan, T. M., & Wolpaw, J. R. (2003). EMG contamination of EEG: Spectral and topographical characteristics. Clinical Neurophysiology, 114 (9), 1580–1593. 10.1016/S1388-2457(03)00093-2

Hyvärinen, A., & Oja, E. (2000). Independent component analysis: Algorithms and applications. Neural Networks, 13 (4–5), 411–430. 10.1016/S0893-6080(00)00026-5

Leys, C., Ley, C., Klein, O., Bernard, P., & Licata, L. (2013). Detecting outliers: Do not use standard deviation around the mean, use absolute deviation around the median. Journal of Experimental Social Psychology, 49 (4), 764–766. 10.1016/j.jesp.2013.03.013

Mognon, A., Jovicich, J., Bruzzone, L., & Buiatti, M. (2011). ADJUST: An automatic EEG artifact detector based on the joint use of spatial and temporal features. Psychophysiology, 48 (2), 229–240. 10.1111/j.1469-8986.2010.01061.x

Nolan, H., Whelan, R., & Reilly, R. B. (2010). FASTER: Fully automated statistical thresholding for EEG artifact rejection. Journal of Neuroscience Methods, 192 (1), 152–162. 10.1016/j.jneumeth.2010.07.015

Oostenveld, R., Fries, P., Maris, E., & Schoffelen, J.-M. (2011). FieldTrip: Open source software for advanced analysis of MEG, EEG, and invasive electrophysiological data. Computational Intelligence and Neuroscience, 2011, 156869. 10.1155/2011/156869

Pion-Tonachini, L., Kreutz-Delgado, K., & Makeig, S. (2019). ICLabel: An automated electroencephalographic independent component classifier, dataset, and website. NeuroImage, 198, 181–197. 10.1016/j.neuroimage.2019.05.026

Rogasch, N. C., Sullivan, C., Thomson, R. H., Rose, N. S., Bailey, N. W., Fitzgerald, P. B., Farzan, F., & Hernandez-Pavon, J. C. (2017). Analysing concurrent transcranial magnetic stimulation and electroencephalographic data: A review and introduction to the open-source TESA toolbox. NeuroImage, 147, 934–951. 10.1016/j.neuroimage.2016.10.031

Rogasch, N. C., Thomson, R. H., Farzan, F., Fitzgibbon, B. M., Bailey, N. W., Hernandez-Pavon, J. C., Daskalakis, Z. J., & Fitzgerald, P. B. (2014). Removing artefacts from TMS-EEG recordings using independent component analysis: Importance for assessing prefrontal and motor cortex network properties. NeuroImage, 101, 425–439. 10.1016/j.neuroimage.2014.07.037

Ryali, S., Glover, G. H., Chang, C., & Menon, V. (2009). Development, validation, and comparison of ICA-based gradient artifact reduction algorithms for simultaneous EEG-spiral in/out and echo-planar fMRI recordings. NeuroImage, 48 (2), 348–361. 10.1016/j.neuroimage.2009.06.072

Tamburro, G., Croce, P., Zappasodi, F., & Comani, S. (2021). Automated detection and removal of cardiac and pulse interferences from neonatal EEG signals. Sensors, 21 (19), 6364. 10.3390/s21196364

Veniero, D., Bortoletto, M., & Miniussi, C. (2009). TMS-EEG co-registration: On TMS-induced artifact. Clinical Neurophysiology, 120 (7), 1392–1399. 10.1016/j.clinph.2009.04.023

Winkler, I., Haufe, S., & Tangermann, M. (2011). Automatic classification of artifactual ICA-components for artifact removal in EEG signals. Behavioral and Brain Functions, 7, 30. 10.1186/1744-9081-7-30

Wu, W., Keller, C. J., Rogasch, N. C., Longwell, P., Shpigel, E., Rolle, C. E., & Etkin, A. (2018). ARTIST: A fully automated artifact rejection algorithm for single-pulse TMS-EEG data. Human Brain Mapping, 39 (4), 1607–1625. 10.1002/hbm.23938

